# Modeling circadian rhythm splitting under chronic light and the asymmetric transition kinetics between lighting conditions

**DOI:** 10.64898/2026.11.04.764283

**Authors:** Alexander V. Goltsev, José F. F. Mendes

## Abstract

In various vertebrates, chronic light splits the single daily 24-h circadian activity bout into two 12-h behavioral components per cycle, yet the underlying mechanism remains unresolved. Here, we develop a two-lobe biophysical model of the suprachiasmatic nucleus (SCN) to understand this phenomenon. By representing the master clock as interacting ventral core and dorsal shell populations of Kuramoto-type oscillators across bilateral lobes, we suggest that chronic light changes the excitation/inhibition (E/I) balance orchestrated by GABA, which either inhibits or excites SCN neurons. Under constant darkness, excitation dominates and provides in-phase synchronization of the populations. Under chronic light, inhibition dominates, and the ipsilateral and contralateral couplings among the core and shell populations become negative. Consequently, the shells in the left and right lobes oscillate in antiphase, producing the 12-h behavioral rhythm in downstream tissues. We suggest that this E/I balance is shifted by a light-induced change in the expression of chloride cotransporters in SCN neurons. Another key novelty is the introduction of the biologically motivated dynamics of chloride cotransporter expression, which explains the sign inversion of ipsilateral and contralateral couplings under time-varying lighting conditions. The model also reproduces the effects of unilateral SCN lesions: if one lobe is abolished, the remaining intact lobe autonomously stabilizes into a single-bout circadian output with a period comparable to that of the intact system. Finally, we compare our results with experimental data and demonstrate a close agreement.

## Introduction

Exposing animals to constant light (LL) for 30–110 days can induce circadian rhythm ‘splitting’, wherein the single daily activity bout bifurcates into two components roughly 12 hours apart. Observed across diverse vertebrates—including rodents, primates, tree shrews, starlings, and lizards—splitting causes behavioral, thermal, and endocrine outputs to peak twice daily; see (Pittendrigh (1960); Pittendrigh and Dann (1976); Pickard et al. (1984); Swann and Turek (1985); Yan et al. (2005); Butler et al. (2012); Evans and Gorman (2016)) and references therein. In split animals, chronic light changes relative phases of molecular oscillations in ventral (core) and dorsal (shell) regions of the left and right lobes of the suprachiasmatic nucleus (SCN). When the shell in the right lobe exhibits night-typical activity, the shell in the left lobe displays day-typical activity. Then, 12 h later, the right and left lobes demonstrate the opposite behavior. For example, when the core in the left SCN lobe shows day-typical elevation of Period1 (PER1) expression, the contralateral core in the right SCN lobe shows night-typical elevation of Bmal1 expression (de la Iglesia et al. (2000, 2003); Ohta et al. (2005); Tavakoli-Nezhad and Schwartz (2005); Yan et al. (2005); Butler et al. (2012)). Another interesting feature of the split state was observed in (Yan et al. (2005)), where it was reported that the PER1 rhythms in the ipsilateral core and shell regions of the split animal are also in antiphase within each SCN lobe. Based on these observations and the fact that the SCN shells coordinate behavioral rhythms in downstream tissues throughout the body and brain (Abrahamson and Moore (2001); Evans and Gorman (2016); Michel and Meijer (2020); Hastings et al. (2018); Silver (2018)), it can be concluded that the two peaks in behavioral and physiological circadian rhythms in split animals correspond to two subsequent peaks, 12 h apart, of activity in the shells of the left and right lobes, respectively. In contrast to split animals, molecular rhythms in the contralateral regions of the left and right lobes in unsplit animals are strongly correlated in phase. Molecular rhythms in the shells of the left and right lobes form a single daily bout of activity per 24-h circadian cycle. Interestingly, the molecular rhythms in the SCN shells phase-advance the molecular rhythms in the cores; see, for example, (Taylor et al. (2017); Goltsev et al. (2022)). Contralateral couplings, which extend between the left and right SCN lobes across the midline, play an important role in the LL-induced antiphase oscillations of the left and right lobes. This fact is supported by the observation that an unsplit circadian rhythm is restored by unilateral SCN lesions (Pickard and Turek (1982, 1983)). Unilateral SCN lesions, which destroy more than 50% of a single SCN lobe, restore an unsplit rhythm with a period of approximately 24 h, while lesions destroying less than 50% of a single SCN lobe do not abolish the split condition and have little effect on the split rhythm. The data discussed above indicate that each bout of the split rhythm is generated by a separate SCN lobe, which is capable by itself of maintaining a circadian rhythm with an approximately 24-h period when the opposite lobe is abolished. However, Harrington et al. (1990) have shown that unilateral SCN damage is not the only method for eliminating the split condition, as lesions of non-SCN tissue can also abolish splitting.

Transferring animals with a split activity rhythm from LL to constant darkness (DD) also restores the 24-h unsplit rhythm. This occurs remarkably fast, within 1–4 days (Boulos and Terman (1979); Earnest and Turek (1982)). Subsequent re-exposure to LL after 10–30 days in DD reinduces splitting strikingly rapid, within another 1–4 days, sharply contrasting with the more than 30 days required for de novo splitting in LL (Earnest and Turek (1982)). The mechanism governing these rapid, history-dependent transitions remains unknown.

Despite progress in identifying the neuroanatomical substrates for splitting, the neurophysiological process by which the SCN reorganizes under chronic light remains unclear. It has been proposed that LL-induced splitting may reflect light-induced changes in the nature of coupling among SCN neurons, shifting from excitatory to inhibitory due to alterations in SCN neurophysiology; see (Daan and Berde (1978); Kawato and Suzuki (1980); Oda and Friesen (2002); Meijer et al. (2010, 2012); Evans et al. (2013); Evans and Gorman (2016)). A crucial factor in understanding this coupling shift is that almost all SCN neurons are GABAergic, and the inhibitory transmitter *γ*-aminobutyric acid (GABA) is the most abundant neurotransmitter within the SCN (Abrahamson and Moore (2001)). Acting through A-type receptors, GABA mediates cell-to-cell communication during the daily light-dark (LD) cycle (Liu and Reppert (2000)) and can function either as an inhibitory or an excitatory transmitter; see (Choi et al. (2008); McNeill et al. (2018, 2020)) and references therein. When operating as an excitatory transmitter, GABA couples the dorsal and ventral regions (Liu and Reppert (2000); Shirakawa et al. (2000); Albus et al. (2005)). In the SCN, there are other coupling factors, such as vasoactive intestinal polypeptide (VIP) and arginine vasopressin (AVP), which play an especially important role in the synchronization of neurons in the SCN core and shell, respectively (Evans and Gorman (2016); Aton and Herzog (2005); Boareto et al. (2025); Maywood et al. (2011)). We assume that VIP and AVP preserve synchronization among neurons within the core and shell under chronic light, although direct experimental verification is currently unavailable due to a lack of data. GABAergic polarity depends on the excitation/inhibition (E/I) balance, which is set by cation-chloride cotransporters (CCCs) (Yamada et al. (2004); Choi et al. (2008); Blaesse et al. (2010); Belenky et al. (2010); Kaila et al. (2014); Myung et al. (2015); Sulihu et al. (2021)). Two primary cotransporters maintain intracellular chloride homeostasis: NKCC1 imports chloride, elevating internal concentrations to render GABA excitatory (depolarizing), whereas KCC2 exports chloride, making GABA hyperpolarizing and inhibitory (Tillman et al. (2019)). The expression ratio of KCC2 to NKCC1 thus determines whether net GABA-mediated coupling is excitatory or inhibitory. The excitatory response is typical of neurons with a low ratio of KCC2 to NKCC1 (Choi et al. (2008); Alamilla et al. (2014); Irwin and Allen (2009); McNeill et al. (2018, 2020)). Environmental light directly modulates NKCC1 and KCC2 expression in the SCN (Farajnia et al. (2014); Myung et al. (2015); Engberink et al. (2018)). Modulation of intracellular chloride controls the network encoding of seasonal time, adjusting the strength and polarity of GABA_*A*_-mediated synaptic input (Myung et al. (2015)). However, the precise mechanisms underlying this effect remain to be fully elucidated.

Several biologically motivated models have been proposed to describe splitting (Pittendrigh and Dann (1976); Daan and Berde (1978); Kawato and Suzuki (1980); Turek et al. (1982); Morin and Cummings (1982); Swann and Turek (1985); Lees et al. (1983); Oda and Friesen (2002); Yan et al. (2005); Rohling and Meylahn (2020)); however, the understanding of this phenomenon remains incomplete. In this paper, we develop a two-lobe biophysical model of the SCN to mimic the splitting of the circadian rhythm. This model represents the master clock as interacting ventral core and dorsal shell populations of Kuramoto-type oscillators across bilateral lobes. The novelty of our approach lies in demonstrating that splitting can be explained by assuming that chronic light changes the sign of both ipsilateral and contralateral couplings among these populations from positive to negative. Intra-population couplings among neurons within the cores and shells must remain positive to ensure the collective behavior of these sub-populations as single units. In the split state, the neuronal populations in the left and right lobes oscillate in antiphase, in contrast to their in-phase oscillation in constant darkness. We also show that removing contralateral inhibitory connections abolishes inter-lobe competition, allowing the intact lobe to sustain coherent, single-bout circadian oscillations. Another key novelty is the introduction of biologically motivated dynamics of chloride cotransporter expression, which explains the sign inversion of ipsilateral and contralateral couplings under time-varying lighting conditions. Finally, we compare our results to experimental data.

## Methods

In this section, we present a two-lobe biophysical model to describe the SCN dynamics and analyze the emergence of the split circadian rhythm.

### Two-lobe SCN model

Our treatment of splitting employs a biologically motivated core-shell model, which describes mammalian SCN dynamics at the population level in both steady and transient states after a rapid phase shift of the LD cycle, wherein the left and right lobes are synchronized (Goltsev et al. (2022); Aleixo et al. (2025)). The two-lobe model takes into account that: (i) SCN neurons are weakly interacting autonomous oscillators that continue to express endogenous circadian rhythms even when synaptic communication is blocked Liu et al. (1997); Welsh et al. (2010); (ii) the free-running periods of the endogenous oscillators in the core and shell are heterogeneous; (iii) the SCN network exhibits a hierarchical modular organization, i.e., the division of the SCN into two lobes composed of core (ventral) and shell (dorsal) regions (Moore et al. (2002)); (iv) intra-population coupling among oscillators within the cores and shells; (v) ipsilateral and contralateral couplings between the core and shell populations within and across the lobes; and (vi) the photic cue acts on the core oscillators. We aim to generalize this core-shell model to account for splitting, wherein the two interacting lobes oscillate in antiphase.

The two-lobe SCN model is schematically represented in Figure 1. There are four populations of neurons numbered by the index *n* = 1, 2, …, 4. Populations 1 and 2 represent the core and shell in the left SCN lobe, respectively, while populations 3 and 4 represent the core and shell in the right SCN lobe. Neurons within population *n* are mutually coupled with strength *K*_*nn*_. The core and shell neurons are connected by the couplings *K*_12_ and *K*_21_ in the left lobe, and by *K*_34_ and *K*_43_ in the right lobe. The neurons in cores 1 and 3 are coupled to each other by the contralateral couplings *K*_13_ and *K*_31_, whereas the neurons in shells 2 and 4 are coupled by the contralateral couplings *K*_24_ and *K*_42_. Commissural couplings between populations 1 and 4, as well as populations 2 and 3, are absent in our model because they have not been observed in neuroanatomical investigations (Moore et al. (2002)).

**Figure 1.**
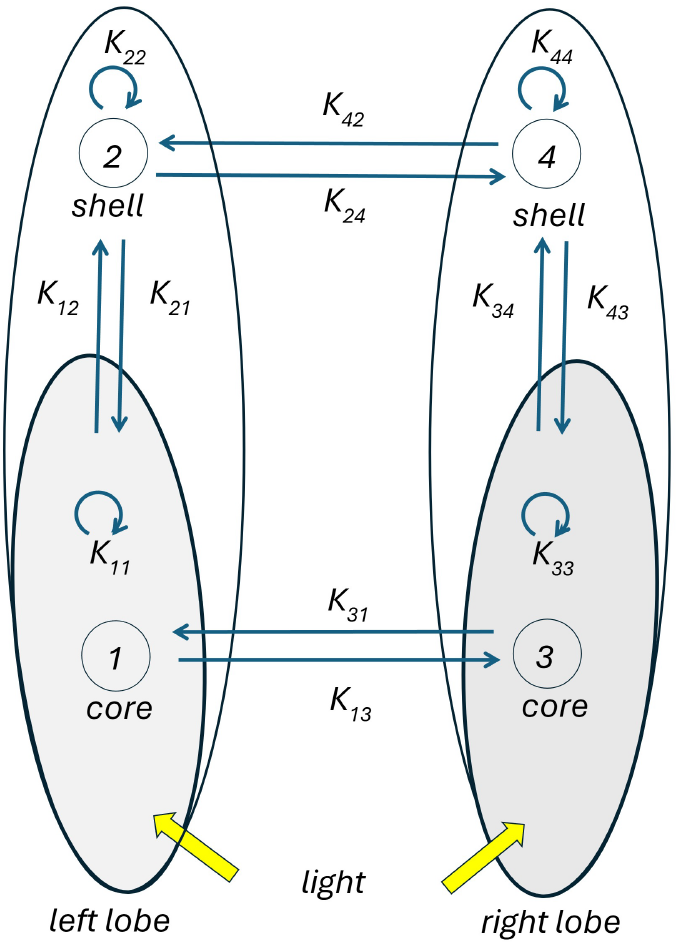
Schematic representation of the two-lobe model consisting of four populations of endogenous oscillators in the SCN. Circles 1 and 2 represent the core and shell in the left lobe of the SCN, respectively. Circles 3 and 4 represent the core and shell in the right lobe. *K*_11_, *K*_22_, *K*_33_, and *K*_44_ are intra-couplings among oscillators within the populations 1, 2, 3, and 4. *K*_12_, *K*_21_, *K*_34_, and *K*_43_ are ipsilateral couplings in the left and right lobes. *K*_13_, *K*_31_, *K*_24_ and *K*_42_ are the contralateral couplings. The yellow arrows represent chronic light acting on oscillators in cores 1 and 3.

We assume that the dynamics of these autonomous oscillators, communicating via multiple spikes throughout the circadian cycle, can be accurately approximated by a set of phase equations that depend only on the phase differences between the oscillators (Kuramoto (1975); Ermentrout and Kopell (1991)). Within this approximation, a positive coupling constant between oscillators results from excitatory signaling from one clock cell to another, while a negative coupling constant corresponds to inhibitory signaling. At the population level, within the two-lobe SCN model, the dynamical state of the core and shell is described by eight variables: *ρ*_*n*_ and *ψ*_*n*_, where *n* = 1, 2, …, 4. The synchronization index *ρ*_*n*_ characterizes the extent of phase alignment between the oscillators within population *n* and varies between 0 and 1. When *ρ*_*n*_ = 0, the oscillators within population *n* are in an asynchronous state, whereas *ρ*_*n*_ = 1 corresponds to a fully synchronized state. The group phase *ψ*_*n*_ represents the average phase of population *n* in the synchronized state.

Given an initial state at *t* = 0, the synchronization indices *ρ*_*n*_(*t*) and the group phases *ψ*_*n*_(*t*) evolve according to eight coupled nonlinear dynamical equations:

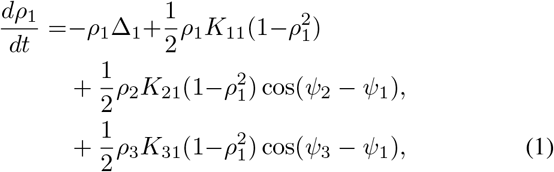

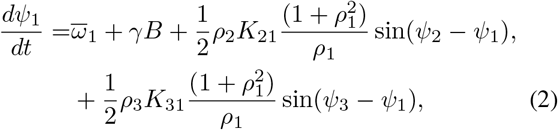

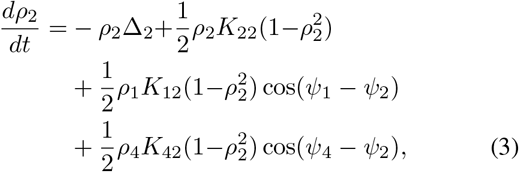

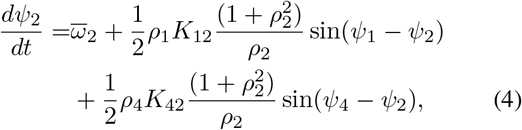

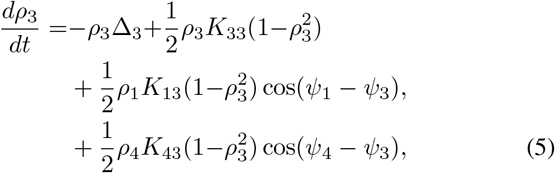

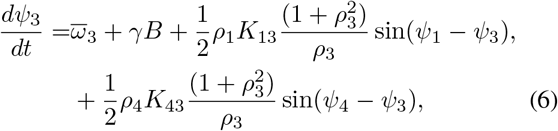

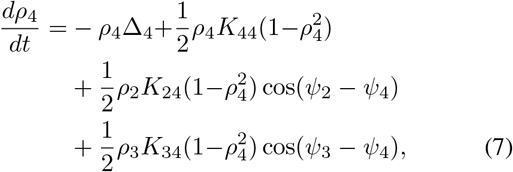

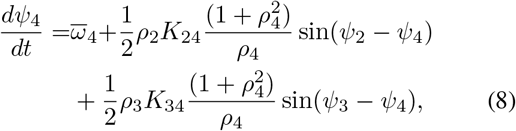

where 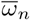 is the average group frequency of autonomous oscillators in populations *n, n* = 1, 2, …, 4. The parameter Δ_*n*_ denotes the spread of the natural frequency distribution of the free-running frequencies of the oscillators in the population *n*. The phenomenological parameter *γB* can be positive or negative. The parameter *γ* characterizes the susceptibility of the core phases to the photic cue with intensity *B*. In general, the parameter may depend on time (Taylor (2021); Aleixo et al. (2025)). The sign of *γB* is chosen to satisfy Aschoff’s first rule (Aschoff (1979); Goltsev et al. (2022); Aleixo et al. (2025)). The parameter *γB* is negative for nocturnal animals and positive for diurnal animals. Note that Eqs. (1)-(8) are derived assuming that light acts on all core cells within the bilateral SCN. In vivo, only about 75% of core cells receive direct retinal innervation (Meijer et al. (1986)). However, this is enough to entrain the entire population of oscillators (Yoon et al. (2021)). Note that according to Eqs. (1)–(8), synchronization *ρ*_*n*_ among oscillators in a population *n* can vary in time. It is important that if the intracoupling *K*_*nn*_ is either negative or positive but smaller than the critical threshold 2Δ_*n*_ and couplings with other compartments are absent, then the oscillators fail to synchronize due to fluctuations breaking down collective synchronization of a many-oscillators system (Goltsev et al. (2022)), although a two-oscillators system with negative coupling can be synchronized. Thus, according to the phase transition theory, to provide collective synchronization between oscillators within each SCN core and shell, the intracouplings must be positive and sufficiently large.

### Steady state of the two-lobe SCN model

Let us find the steady-state solutions of Eqs. (1) to (8) for the case where the parameters of the left and right lobes are identical. We also assume that the left and right cores—populations 1 and 3, respectively—receive identical photic input. This symmetric configuration corresponds to the following relations among the model parameters:

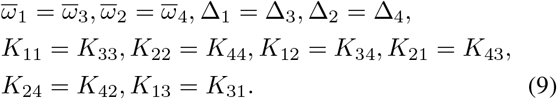

We are looking for a steady state solution of Eqs. (1)-(8), assuming that the SCN compartments oscillate with the common circadian frequency *ω*_*LL*_. In the steady state, we have *dρ*_*n*_*/dt* = 0 and *dψ*_*n*_*/dt* = *ω*_*LL*_, therefore

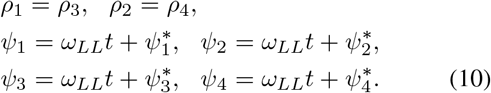

Here 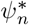 is the group phase of the compartment *n* at *t* = 0. Solving Eqs. (1) to (8), we find that the circadian frequency *ω*_*LL*_ equals

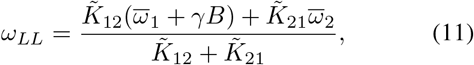

where 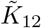 and 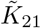 are the effective couplings:

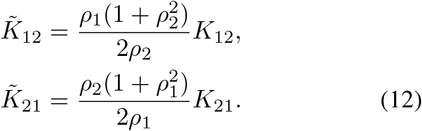

The phase differences, *ψ*_2_ − *ψ*_1_ and *ψ*_4_ − *ψ*_3_, between the group phases of the shells and cores of the left and right lobes, are given by the following equation:

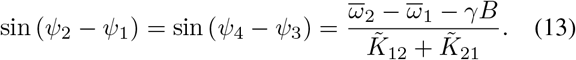

If all couplings among the SCN neurons are positive, then under DD conditions, i.e., at *γB* = 0, the SCN exhibits an unsplit circadian rhythm. The corresponding stability condition is given by:

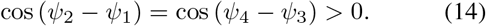

The group phases of the cores and shells in the left and right lobes are the same, *ψ*_1_ = *ψ*_3_ and *ψ*_2_ = *ψ*_4_. In this case, the equation (13) gives the phase angle difference:

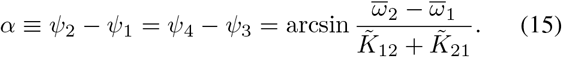

Therefore, the shells lead the cores. Under DD, the circadian frequency is

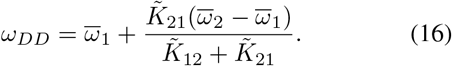

We assume that under the chronic light condition, the intra-population couplings remain positive, whereas the ipsilateral and contralateral couplings among the SCN compartments become negative:

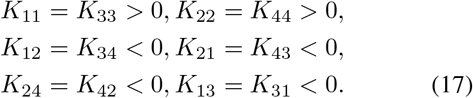

The steady state solution of Eqs. (1) to (8) is stable at the following conditions:

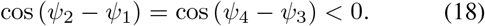

The core and shell in the left lobe oscillate in antiphase to the core and shell in the right lobe:

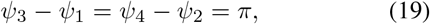

The fact that the compartments of the left and right lobes oscillate in antiphase means that in the split state, the SCN shells send signals to downstream tissues twice per circadian cycle *τ*_*LL*_. This directly explains the splitting of the behavioral rhythm into two bouts, as observed experimentally.

According to Eqs. (13) and (18), the phase angle difference between the core and shell in the left and right lobes is

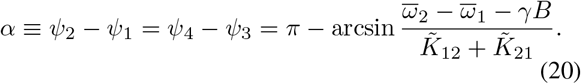

If 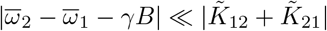, then *ψ*_2_ − *ψ*_1_ = *ψ*_4_ − *ψ*_3_ ≈ *π*, i.e., in each lobe the core and shell oscillate almost in antiphase, in agreement with empirical findings (de la Iglesia et al. (2000, 2003); Ohta et al. (2005); Yan et al. (2005)). Comparing Eq. (15) to Eq. (20) reveals that transferring from DD to LL increases the phase angle difference *α* = *ψ*_2_ − *ψ*_1_ = *ψ*_4_ − *ψ*_3_ between the core and shell in the left and right lobes, see Figure 2.

**Figure 2.**
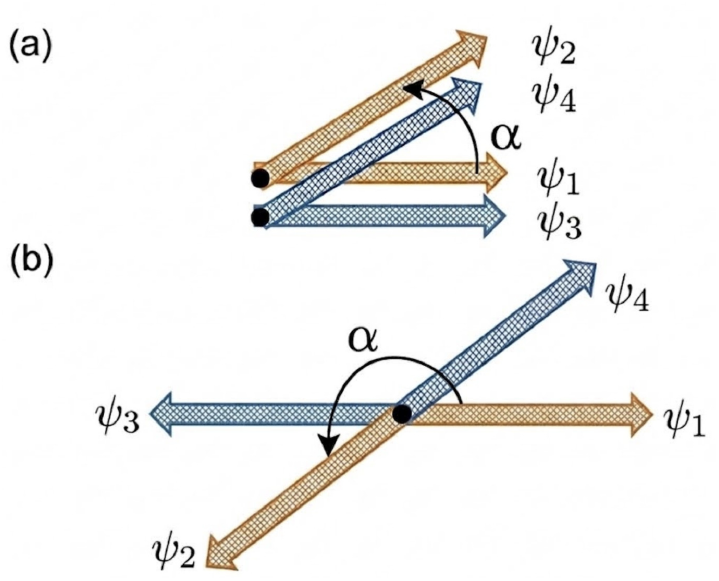
Relative group phases of the cores and shells in the left and right lobes in unsplit (a) and split (b) steady states of the SCN, see Fig. 1. The parameter *α* = *ψ*_2_ *− ψ*_1_ defines the stable phase angle difference between the group phases of the shell and core.

## Results

Within the two-lobe model of the SCN we consider the following cases: (1) unsplit rhythm under DD conditions, when all couplings are positive; (2) split rhythm under LL conditions, when ipsilateral and contralateral couplings are negative; (3) rhythm after unilateral lesions under DD and LL conditions. For these purposes, we implement a fourth-order Runge-Kutta numerical integration method to evaluate the full eight-dimensional coupled population model, see Eqs. (1) to (8). The model parameters are listed in Table 1. They are taken from experimental data on the SCN of mice at 12:12 h LD cycles (Goltsev et al. (2022); Aleixo et al. (2025); Taylor et al. (2017)). In numerical calculations of Eqs. (1) to (8), we use dimensionless parameters by rescaling all parameters with respect to the core frequency unit Δ_1_: couplings, *K*_*nm*_*/*Δ_1_→*K*_*nm*_; time, *t*Δ_1_→*t*; the optic cue, *γB/*Δ_1_→*b* = *B/B*_0_, where *B*_0_ is a characteristic unit of the photic cue intensity depending on the type of animal. We use a dimensionless value *γB/*Δ_1_ = −0.5 that corresponds to mice under the constant light 10 lux (Goltsev et al. (2022)). Based on the anatomical organization of the SCN (Moore et al. (2002)), we assume that the strengths of contralateral couplings *K*_13_ and *K*_24_ are smaller than those of the ipsilateral couplings *K*_12_ and *K*_34_. Albus et al. (2005) measured the electrical activity of the unsplit SCN in rats and observed that the ventral part produces a stronger phase-shifting effect on the dorsal part than vice versa. This effect can be interpreted as indicating that signaling from the core to the shell is stronger than the reciprocal signaling from the shell to the core, i.e., |*K*_12_| *>* |*K*_21_| and |*K*_34_| *>* |*K*_43_|.

**Table 1.**
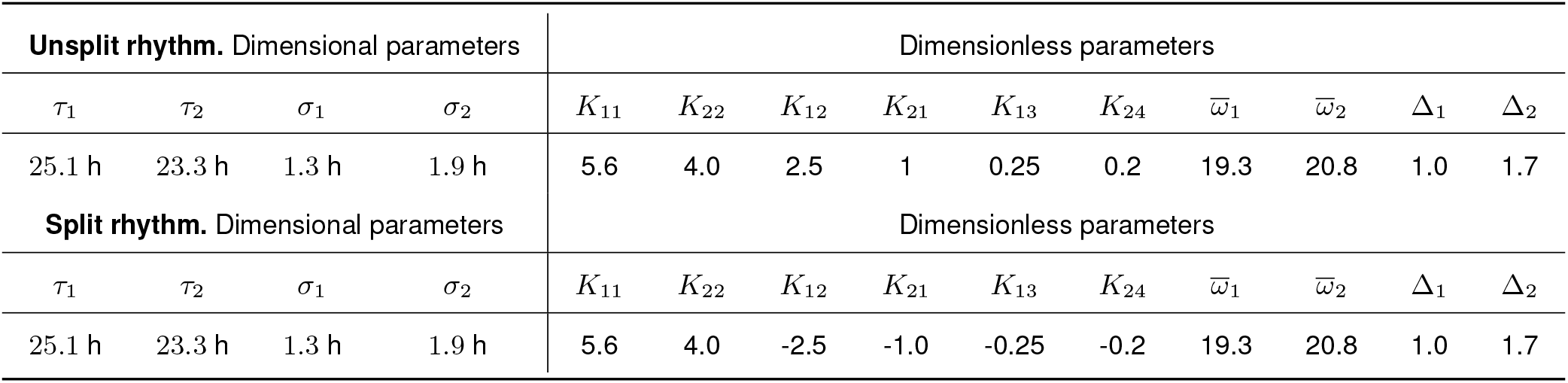
List of parameters in Eqs. (1)–(8) for the two-lobe SCN model with identical lobes, Eq. (9): *τ*_1(2)_ represent the average free-running period of the core (shell); *σ*_1(2)_ denotes the standard deviation of free-running periods; *K*_11_ and *K*_22_ specify the intracoupling in the core and shell; *K*_12_ and *K*_21_ indicate the core–shell ipsilateral couplings; *K*_13_ and *K*_24_ indicate the core–core and shell–shell contralateral couplings; 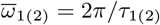 defines the average free-running frequency of core (shell) oscillators; 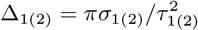 represents the Lorentzian spread of free-running frequencies in the core (shell). We use dimensionless value *γB→−*0.5 that corresponds to mice under the constant light 10 lux (Goltsev et al. (2022)).

| **Unsplit rhythm.** Dimensional parameters |  |  |  | Dimensionless parameters |  |  |  |  |  |  |  |  |  |
|---|---|---|---|---|---|---|---|---|---|---|---|---|---|
| τ<sub>1</sub> | τ<sub>2</sub> | σ<sub>1</sub> | σ<sub>2</sub> | K<sub>11</sub> | K<sub>22</sub> | K<sub>12</sub> | K<sub>21</sub> | K<sub>13</sub> | K<sub>24</sub> | ω̄<sub>1</sub> | ω̄<sub>2</sub> | Δ<sub>1</sub> | Δ<sub>2</sub> |
| 25.1 h | 23.3 h | 1.3 h | 1.9 h | 5.6 | 4.0 | 2.5 | 1 | 0.25 | 0.2 | 19.3 | 20.8 | 1.0 | 1.7 |
| **Split rhythm.** Dimensional parameters |  |  |  | Dimensionless parameters |  |  |  |  |  |  |  |  |  |
| τ<sub>1</sub> | τ<sub>2</sub> | σ<sub>1</sub> | σ<sub>2</sub> | K<sub>11</sub> | K<sub>22</sub> | K<sub>12</sub> | K<sub>21</sub> | K<sub>13</sub> | K<sub>24</sub> | ω̄<sub>1</sub> | ω̄<sub>2</sub> | Δ<sub>1</sub> | Δ<sub>2</sub> |
| 25.1 h | 23.3 h | 1.3 h | 1.9 h | 5.6 | 4.0 | -2.5 | -1.0 | -0.25 | -0.2 | 19.3 | 20.8 | 1.0 | 1.7 |

### Unsplit circadian rhythm under DD conditions

We solved numerically Eqs. (1) to (8) under DD, i.e., *γB* = 0, using parameters in Table 1. In the steady state, we find the core and shell synchronization indices: *ρ*_1_ = *ρ*_3_ ≈ 0.88 and *ρ*_2_ = *ρ*_4_ ≈ 0.65. Equations (12) to (15) give the effective scaled couplings and the phase angle difference: 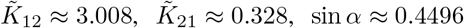. The stable phase angle difference is *α* ≈ 0.466 rad (26.7^*°*^). Because *α* = *ψ*_2_ − *ψ*_1_ = *ψ*_4_ − *ψ*_3_ *>* 0, the shell group phase leads that of the cores, see Fig. 1a, which matches the baseline biological observations. The DD circadian period is *τ*_*DD*_ ≈ 24.96 h in agreement with experimental observations in rats. Campuzano et al. (1998) reported *τ*_*DD*_ = 24.5 h. Under DD, the activities of the left and right shells oscillate with the common frequency and phase, *ψ*_2_(*t*) = *ψ*_4_(*t*). They provide outputs to downstream tissues with the period *τ*_*DD*_.

### Split circadian rhythm under LL conditions

To model splitting, we numerically solve Eqs. (1) to (8) assuming that chronic light changes the sign of ipsilateral and contralateral couplings from positive to negative, see Table Under chronic light conditions with photic drive *γB* = −0.5, internal synchronization of core and shell populations remains strong due to strong positive internal couplings (*K*_11_ = 5.6, *K*_22_ = 4.0). The synchronization indices are *ρ*_1_ = *ρ*_3_ ≈ 0.84 and *ρ*_2_ = *ρ*_4_ ≈ 0.62, which are slightly lower than the values 0.88 and 0.65 in the unsplit state. Using Eqs. (12) and (13) yields: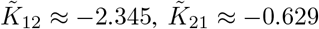, sin *α* ≈ −0.5044. Applying the stability condition Eq. (18), we find the phase angle difference,

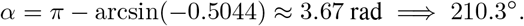

Moreover, in the split state we have *ψ*_3_ = 180^*°*^+ *ψ*_1_ and *ψ*_4_ = 180^*°*^+ *ψ*_2_. This implies that the core and shell in the right lobe oscillate in strict antiphase to their contralateral counterparts. The phase relationships between group phases of the cores and shells in the split state are shown in Fig. 2. Thus, the underlying property of the split state is that the core and shell within each individual lobe are driven into a near-antiphase relationship (210.3^*°*^≈ 180^*°*^+ 30.3^*°*^) and the populations of the left and right lobes oscillate in antiphase with equal frequencies in close agreement with experimental findings (de la Iglesia et al. (2000, 2003); Ohta et al. (2005); Tavakoli-Nezhad and Schwartz (2005); Yan et al. (2005); Butler et al. (2012)). In the split state, the period is *τ*_*LL*_ = 24.75 h. A comparison of the circadian periods under LL and DD conditions, *τ*_*LL*_ = 24.75 h and *τ*_*DD*_ = 24.96 h, reveals that each SCN lobe accelerates its pace when split. This matches the known behavioral transition where transferring split animals back to darkness causes their active clock period to lengthen. Our estimation of the period truncation, *τ*_*DD*_ − *τ*_*LL*_ ≈ 0.21 h, demonstrates excellent agreement with empirical reports, see Table 1 in (Earnest and Turek (1982)).

To elucidate why under chronic light the single daily behavioral activity bout bifurcates into two strict antiphase components, we note that the SCN shells provide the primary outputs to downstream tissues, thereby driving behavioral rhythms (Abrahamson and Moore (2001); Evans and Gorman (2016); Michel and Meijer (2020); Hastings et al. (2018); Silver (2018)). As we showed above, under constant lighting, the left and right shells oscillate in strict antiphase with the period *τ*_*LL*_ = 24.75 h. Because the left and right shells project to downstream tissues, their combined signaling produces the split behavioral rhythm with a half-period of *τ*_*LL*_*/*2 ≈ 12 h. In Figure 3b, we plot the outputs of the left and right lobes, *g*_2_(*t*) and *g*_4_(*t*), respectively. For simplicity we use the following function:

**Figure 3.**
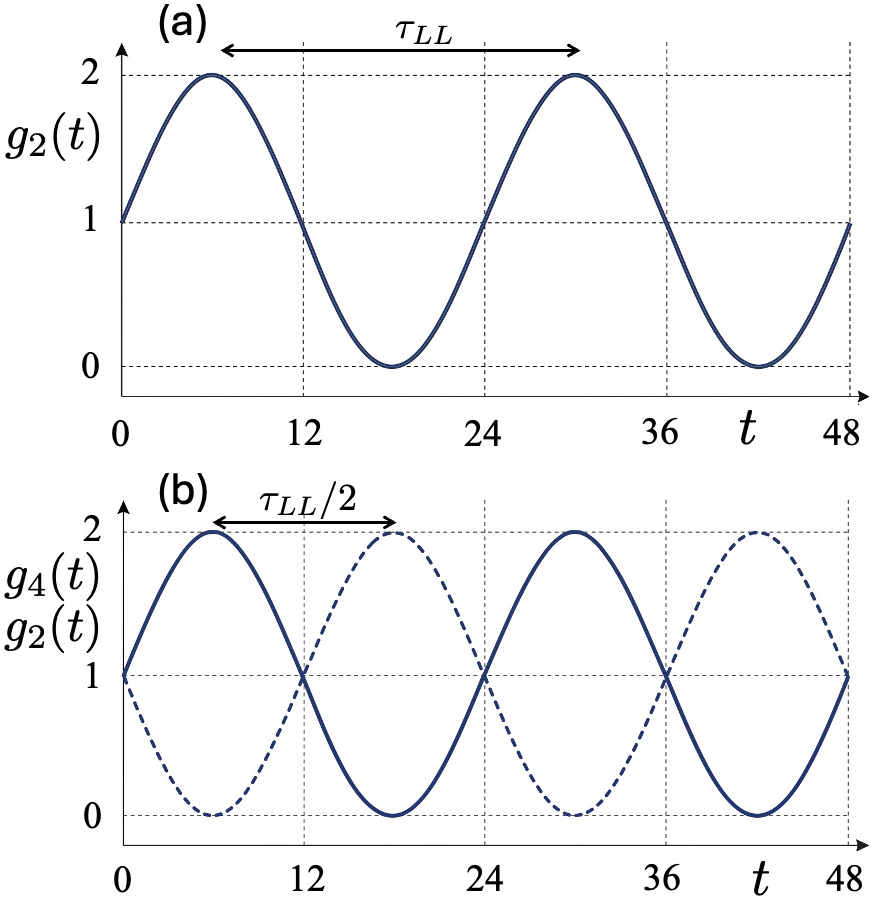
Outputs of the left and right shells: *g*_2_(*t*) and *g*_4_(*t*), solid and dashed curves, respectively. (a) Under DD, we have *g*_2_(*t*) = *g*_4_(*t*). (b) Under LL, the left and right shells oscillate in antiphase with a periodicity *τ*_*LL*_. The combined signaling produced by the shells reaches a maximum with a periodicity *τ*_*LL*_*/*2. It leads to a split behavioral rhythm in the downstream tissues.

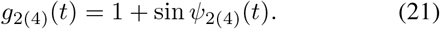

Choosing an alternative output function does not alter the primary conclusion that under chronic light the combined signaling produced by the left and right lobes oscillates at twice the frequency of the unsplit circadian rhythm.

### Unilateral lesion of the split SCN

Experimentally, the effect of a unilateral SCN lesion of the bilaterally paired SCN lobes on the split circadian rhythm was studied in (Pickard and Turek (1982, 1983)). In this section we model unilateral lesion by removing the right lobe in the set of equations Eqs. (1)-(8) with negative ipsilateral and contralateral couplings. The left lobe is described by Eqs. (1)-(4) where we put *ρ*_3_ = *ρ*_4_ = 0. The ipsilateral couplings *K*_12_ and *K*_21_ are negative and are given in Table 1. An active nocturnal photic driving force is *γB* = −0.5. Numerical computing Eqs. (1)-(4) shows that the lesion causes a slight drop in population synchronization parameters of the left core and shell to *ρ*_1_ ≈ 0.835 and *ρ*_2_ ≈ 0.612. The phase angle difference is *α* ≈ 3.88 rad =⇒ 222.4^*°*^, which is larger than 210.3^*°*^in the intact split rhythm. The corresponding circadian rhythm in the left lobe is *τ*_*LL*_ = 25.26 h. It is slightly longer than the period *τ*_*LL*_ = 24.75 h found above in the split state of the intact SCN. Experimentally, Pickard and Turek (1983) observed animals with shortening and lengthening *τ*_*LL*_ after surgery. The mean change in the period was about 0.36 h which is close to our result Δ*τ* = 0.51 h.

### Unilateral lesion of the SCN under DD

Let us study how a unilateral lesion of bilaterally paired lobes affects the circadian rhythm under the DD condition studied above. We use the positive couplings in Table 1. Eliminating the cross-midline reinforcement terms (*K*_13_ = *K*_24_ = 0) in Eqs. (1)-(8) causes a slight drop in the internal synchronization of the isolated populations: *ρ*_1_ ≈ 0.811 and *ρ*_2_ ≈ 0.605. Recalculating the effective coupling terms gives: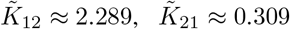, and sin *α* ≈ 0.5774. The resulting phase angle difference shifts to *α* = arcsin(0.5774) ≈ 0.615 rad (35.24^*°*^), and the circadian period after ablation is *τ*_*DD*_ = 24.93 h.

Thus, under DD, ablation shifts the shell-to-core phase angle difference to *α* = 35.24^*°*^and settles the circadian period at *τ*_*DD*_ = 24.93 h, reflecting a minor change (Δ*τ*_*DD*_ = −0.03 h) compared to the intact system. This confirms that an isolated lobe sustains a coherent, single-bout rhythm, aligning with empirical observations where unilateral ablation abolishes splitting (Pickard and Turek (1983); Pickard et al. (1984)). Notably, the intact DD phase angle difference (*α* = 35.24^*°*^) contrasts sharply with the near-antiphase state (*α* = 222.4^*°*^) maintained under persistent negative ipsilateral coupling in LL. Our numerical results are summarized in Table 2.

**Table 2.** Circadian periods computed using the two-lobe model under LL and DD conditions, *τ*_*LL*_ and *τ*_*DD*_, respectively. The data in the ‘intact’ column present results for the intact two-lobe model with the parameters from Table 1. The ‘after UL’ column presents results after a unilateral lesion (UL). Δ*τ* represents the change in the circadian period after a unilateral lesion under LL or DD conditions.

|  | intact | after UL | Δτ |
|---|---|---|---|
| τ<sub>LL</sub> | 24.75 h | 25.26 h | 0.51 h |
| τ<sub>DD</sub> | 24.96 h | 24.93 h | -0.03 h |
| Experimental data | 24.5 h |  |  |

### Microscopic chloride cotransporter dynamics and macroscopic behavioral rhythmicity

In this section, to elucidate the slow onset of splitting under constant light and the rapid transition to an unsplit rhythm upon transfer to DD conditions observed by Earnest and Turek (1982), we model the time-varying expressions of KCC2 and NKCC1 to mimic their impact on circadian rhythms, as well as on the ipsilateral and contralateral coupling among SCN neurons.

Let us track the chloride cotransporter expression ratio *Z*,

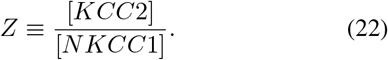

The distance between the cell’s current cotransporter state, *Z*(*t*), and its light-driven target, *Z*_*tg*_, creates a continuous biological driving force that pushes the system to adapt:

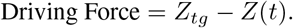

Because synthesizing new cotransporters and restructuring membrane proteins require time, the rate of change *dZ/dt* is constrained by a metabolic time constant *τ*_*Z*_(*B*), which represents the cellular inertia at a given light intensity *B*.

Dividing the driving force by *τ*_*Z*_(*B*) yields the following equation:

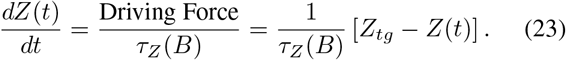

Under DD conditions–namely, at *B* = 0–the system rests at a baseline equilibrium *Z*_0_ where NKCC1 dominates *Z*_0_ *<* 1, maintaining a high intracellular chloride concentration that corresponds to excitatory GABA signaling (Yamada et al. (2004); Choi et al. (2008); Belenky et al. (2010). We propose that chronic light exposure *B >* 0 shifts the balance toward a light-driven steady state with a higher chloride cotransporter expression ratio, *Z*_*B*_ *>* 1, corresponding to inhibitory GABA signaling. We link the evolving cotransporter ratio, *Z*(*t*), directly to our core-shell network coupling coefficients using a continuous hyperbolic tangent switching function:

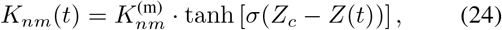

for both ipsilateral and contralateral couplings, as illustrated in Figure 1. Here, 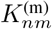 represents the maximum coupling strength, while the parameter *σ* regulates the transition slope from positive to negative coupling. We choose a critical parameter value of *Z*_*c*_ = 1. Therefore, when *Z*(*t*) *<* 1, the couplings *K*_*nm*_(*t*) remain positive, whereas for *Z*(*t*) *>* 1, they become negative. Choosing a different value for *Z*_*c*_ does not alter these qualitative results.

We examine SCN behavior under the time-varying lighting conditions established by Earnest and Turek (1982). Prior to the initial time *t* = 0, the SCN is under DD conditions, and the expression ratio *Z* is equal to its steady-state value, *Z*_0_. The light turns on at time *t* = 0, is turned off at *t*_1_, and is turned on again at *t*_2_, defining three distinct time intervals:

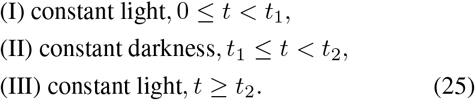

Within this protocol, during time interval I—following the onset of light at *t* = 0—the ratio *Z* increases from the initial condition *Z*(0) = *Z*_0_, tending toward the target value *Z*_*tg*_ = *Z*_*B*_. Under these conditions, Eq. (23) takes the form:

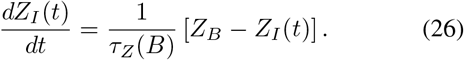

Integrating this equation yields the explicit step-response over time:

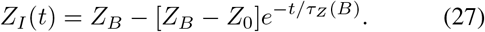

Splitting occurs when the ratio *Z*_*I*_ (*t*) becomes larger than the threshold value of 1, and the couplings in Eq. (24) change sign from positive to negative. This transition takes place at time *t*_*c*1_, which is determined by the following relation:

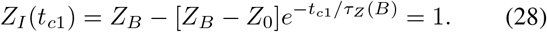

Thus, the circadian rhythm splits after *t*_*c*1_ days under LL conditions. A numerical estimation for *t*_*c*1_ is provided below:

Within the protocol (25), we assume that after 30 days in the split state under LL conditions—that is, at time *t*_1_ = *t*_*c*1_ + 30—the light is turned off. For *t > t*_1_, the SCN is under DD conditions, where the target parameter *Z*_*tg*_ is equal to *Z*_0_ in Eq. (23). Consequently, the ratio *Z*(*t*) decreases according to the following equation:

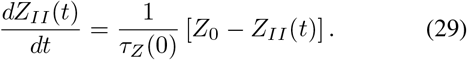

The initial condition for this equation is *Z*_*II*_ (*t* = *t*_1_) = *Z*_*I*_ (*t*_1_). Integrating equation Eq. (29) yields

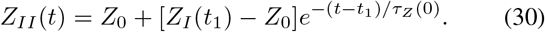

At time *t* = *t*_*c*2_, the chloride cotransporter expression ratio, *Z*_*II*_ (*t*), becomes smaller than the critical value of 1. This corresponds to the transition from the split to the unsplit state. The critical time *t*_*c*2_ is the solution to the following equation:

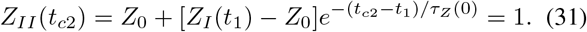

In the interval *t*_*c*1_ *< t < t*_*c*2_, the circadian rhythm remains split because, according to Eq. (24), *Z*_*II*_ (*t*) *>* 1 and the couplings *K*_*nm*_(*t*) are negative. Thus, after the transition from LL to DD, the split state persists for *t*_*c*2_ − *t*_1_ days before transitioning back to the unsplit rhythm.

Following the protocol (25), the light is turned on again at time *t* = *t*_2_. Under LL conditions, the chloride cotransporter expression ratio, *Z*(*t*), increases over time according to the following equation:

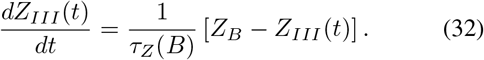

Imposing the initial condition *Z*_*III*_ (*t*_2_) = *Z*_*II*_ (*t*_2_) and integrating this equation yields:

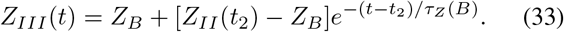

When *Z*_*III*_ (*t*) becomes larger than the critical value of 1, the system transitions from the unsplit to the split circadian rhythm. This transition occurs at time *t* = *t*_*c*3_, which is determined by the following relation:

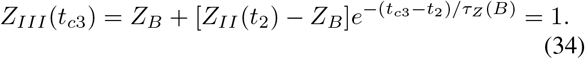

Thus, following the transition from DD to LL, splitting is re-induced after a delay of *t*_*c*3_ − *t*_2_ days.

Following the protocol (25), we numerically solve Eqs. (28), (31), and (34) to mimic the experimental protocol described by Earnest and Turek (1982). The following parameters were used:

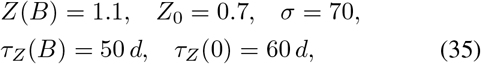

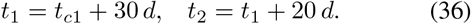

A large value for *σ* in Eq. (24) was chosen to mimic the rapid transition from the unsplit to the split state observed by Earnest and Turek (1982). The large time constants *τ*_*Z*_(*B*) and *τ*_*Z*_(0) describe the slow transitions between the unsplit and split states following changes in lighting conditions. We found that

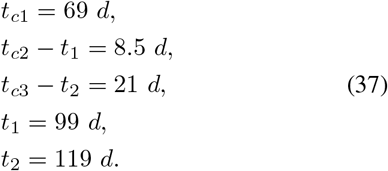

Let us compare the numerical results presented in Fig. 4 with experimental data. First, after transferring the system from DD to LL, the circadian rhythm splits after *t*_*c*1_ = 69 days. This value lies within the experimentally observed range of 30–110 d (Earnest and Turek (1982)). Second, after 30 days under LL conditions, turning off the light abolishes splitting after an interval of *t*_*c*2_ − *t*_1_ = 8.5 days, which is much faster than the 69 days needed to split the rhythm under constant light. Experimental data show a transition of 1–4 days (Earnest and Turek (1982)). Third, we found that subsequent re-exposure to light at *t*_2_ = 119 d re-induces splitting after *t*_*c*3_ − *t*_2_ = 21 days, whereas Earnest and Turek (1982) observed a faster transition of 1–4 days. Although the predicted latencies slightly exceed empirical values, they successfully capture the kinetic asymmetry observed by Earnest and Turek (1982). In the model, this rapid recovery and re-induction stem from three factors: the slow relaxation kinetics of *Z*(*t*), the proximity of the light-adapted steady state *Z*_*B*_ to the bifurcation threshold *Z*_*c*_ = 1, and the fact that the distance from the baseline *Z*_0_ to *Z*_*c*_ is larger than the distance from *Z*_*B*_ to *Z*_*c*_. Consequently, modeling dynamic cotransporter expression satisfactorily accounts for the empirical transition kinetics.

**Figure 4.**
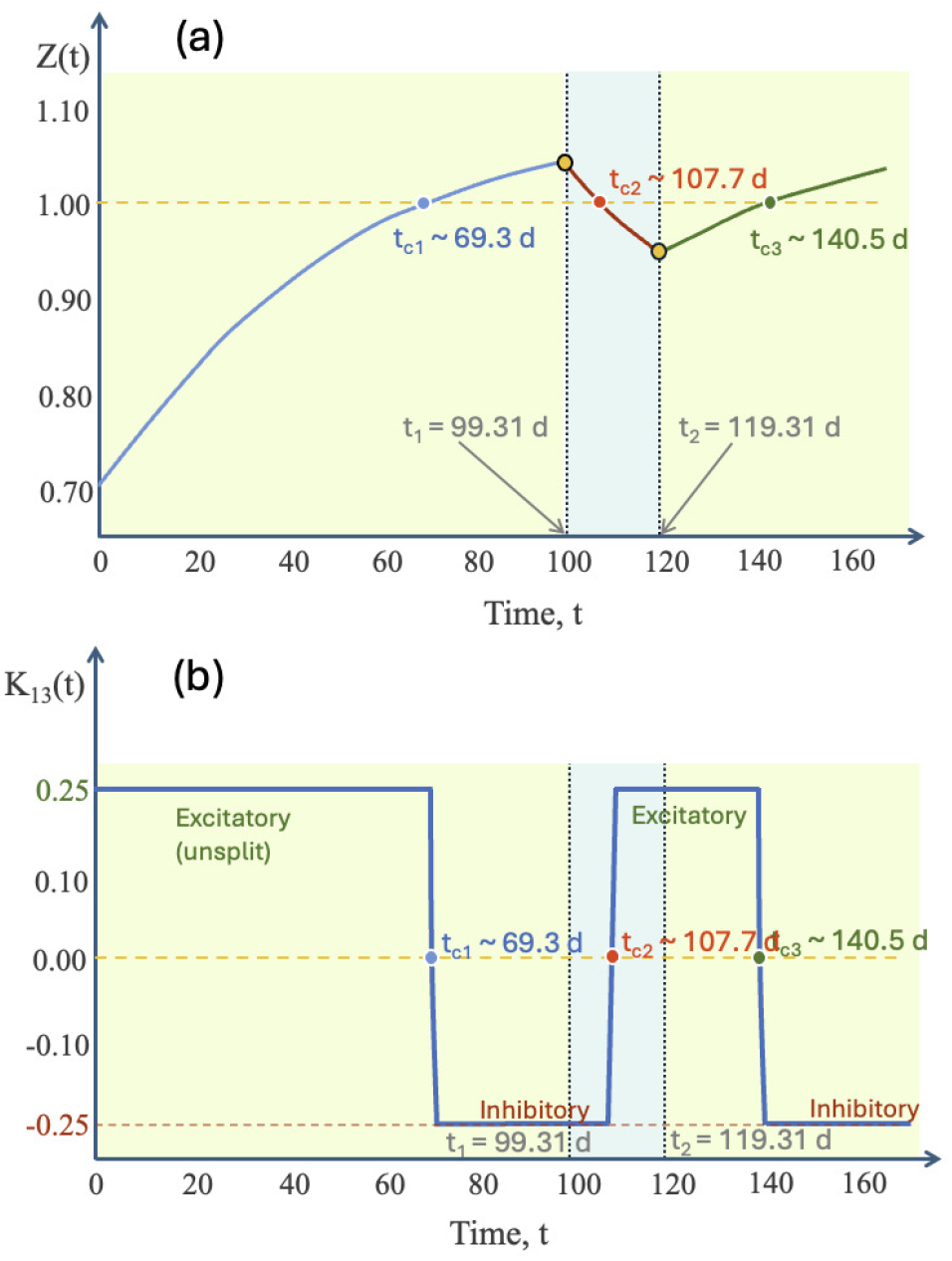
(a) Evolution of the chloride cotransporter expression ratio, *Z*(*t*), across the three time intervals: (I) constant light (0 *≤ t ≤ t*_1_); (II) constant darkness (*t*_1_ *≤ t ≤ t*_2_); and (III) constant light (*t*_2_ *≤ t ≤ t*_3_ + 50). The transition points *t*_*c*1_, *t*_*c*2_, and *t*_*c*3_ correspond to the critical ratio value, *Z* = 1, where the couplings change sign. (b) Evolution of the contralateral coupling coefficient *K*_13_ defined in Eq. (24).

An interesting feature of the splitting process is that, upon transfer from DD to LL, animals maintain an intact circadian rhythm for an extended period exceeding 30 days prior to the rapid onset of splitting. This onset is not characterized by a transient phase during which the two components gradually drift apart. Instead, the two activity components emerge abruptly, approximately 12 hours out of phase with each other (Earnest and Turek (1982)). Figure 4 demonstrates that our model, incorporating a large value for the parameter *σ* in Eq. (24), faithfully mimics this rapid onset of splitting and the absence of a transient phase.

## Conclusions

In this work, a two-lobe biophysical model of the SCN was developed to resolve the mechanisms governing circadian rhythm splitting under chronic light. By representing the mammalian master clock as interacting ventral core and dorsal shell populations of autonomous Kuramoto-type oscillators across bilateral lobes, we link macroscopic behavioral rhythms to microscopic changes in chloride cotransporter dynamics. We suggest that chronic light changes the excitation/inhibition (E/I) balance orchestrated by GABA in SCN neurons. Under dark-dark conditions, excitation dominates and the couplings among the SCN neuronal populations are positive (excitatory). This provides in-phase synchronization of the populations in the left and right lobes. Under chronic light, inhibition dominates and the ipsilateral and contralateral couplings among the core and shell populations become negative (inhibitory), whereas the coupling of neurons belonging to the same population remains positive. We assumed that VIP and AVP preserve synchronization among neurons within the cores and shells under chronic light. These negative couplings stabilize the antiphase oscillations of the cores and shells between the lobes. Because the left and right shells project to downstream tissues 12 h apart, their combined signaling produces the split behavioral rhythm with a half-period of *τ*_*LL*_*/*2 ≈ 12 h.

Based on available experimental data, we proposed that the E/I balance is shifted by the time-varying chloride cotransporter expression ratio *Z*(*t*) = [*KCC*2]*/*[*NKCC*1]. By incorporating cellular metabolic inertia into the time evolution of *Z*(*t*), the model provides a kinetic asymmetry documented experimentally: namely, the protracted induction latency upon initial LL exposure (≈ 69 d), the rapid consolidation upon transfer to DD (≈ 8.5 d), and the accelerated re-emergence of splitting upon subsequent re-exposure to LL (≈ 21 d). These disparate timescales arise naturally from the slow relaxation kinetics of cotransporter expression and the relative proximity of the photic steady-state *Z*_*B*_ to the critical threshold *Z*_*c*_ compared to the dark baseline *Z*_0_. Once the cotransporter ratio *Z*(*t*) crosses the critical threshold, ipsilateral core-shell and contralateral cross-midline couplings reverse their sign from positive to negative. Local intra-compartmental couplings remain positive, preserving coherence within each SCN population. The model also mimics the rapid onset or ending of splitting and the absence of a transient phase when changing lighting conditions from DD to LL or vice versa. The two activity components appear or disappear simultaneously.

Furthermore, the model captures the operational autonomy of single lobes revealed by unilateral SCN ablations. Removing contralateral inhibitory connections abolishes inter-lobe competition, allowing the intact lobe to sustain coherent, single-bout circadian oscillations. While our baseline parameterization predicts a modest period lengthening (Δ*τ* ≈ +0.51 h), empirical studies report both minor shortening (Δ*τ* ≈ −0.17 h) and lengthening (Δ*τ* ≈ +0.28 h) across individual subjects. This indicates that post-lesion pacing is sensitive to subtle differences in residual intra-lobe coupling and intrinsic core-shell frequency detuning.

In conclusion, our findings demonstrate that behavioral splitting is an emergent network property orchestrated by plastic, light-induced polarity shifts in GABAergic signaling. This theoretical framework offers testable predictions regarding the pharmacological and genetic manipulation of chloride cotransporters, KCC2 and NKCC1, in circadian plasticity and provides new insights into multi-oscillator circadian synchronization.

## Acknowledgements

This work was developed within the scope of the project i3N under the project UID/50025/2025 and the Associate Laboratory i3N – LA/P/0037/2020, financed by the FCT.

